# Does early-stage Alzheimer’s disease affect the dynamics of motor adaptation?

**DOI:** 10.1101/2024.01.16.575820

**Authors:** K Sutter, Wijdenes L Oostwoud, RJ van Beers, JAHR Claassen, RPC Kessels, WP Medendorp

**Author notes:** Corresponding author: Prof. dr. Pieter Medendorp.

## Abstract

Alzheimer’s disease (AD) is characterized by an initial decline in declarative memory, while non-declarative memory processing remains relatively intact. Error-based motor adaptation is traditionally seen as a form of non-declarative memory, but recent findings suggest that it involves both fast, declarative and slow, non-declarative adaptive processes. If the declarative memory system shares resources with the fast process in motor adaptation, it can be hypothesized that the fast, but not the slow, process is disturbed in AD patients. To test this, we studied 20 early-stage AD patients and 21 age-matched controls of both sexes using a reach adaptation paradigm that relies on spontaneous recovery after sequential exposure to opposing force fields. Adaptation was measured using error clamps and expressed as an adaptation index (AI). Although patients with AD showed slightly lower adaptation to the force field than the controls, both groups demonstrated effects of spontaneous recovery. The time course of the AI was fitted by a hierarchical Bayesian two-state model in which each dynamic state is characterized by a retention and learning rate. Compared to controls, the retention rate of the fast process was the only parameter that was significantly different (lower) in the AD patients, confirming that the memory of the declarative, fast process is disturbed by AD. The slow adaptive process was virtually unaffected. Since the slow process learns only weakly from error, our results provide neurocomputational evidence for the clinical practice of errorless learning of everyday tasks in people with dementia.

## Introduction

Impaired declarative memory is a hallmark of Alzheimer’s disease (AD), the most common cause of dementia. This progressive memory loss is typically a result of bilateral atrophy of the medial temporal lobe (MTL), including the entorhinal cortex and the hippocampus proper, as well as atrophy in parietal areas (Hyman et al., 1984; Braak and Braak, 1991, 1996; Van Hoesen et al., 1991; Foundas et al., 1997). Despite this loss, there is evidence that aspects of learning and memory that rely more on automatic and unconscious processing, referred to as nondeclarative or procedural memory, are relatively intact (Shadmehr et al., 1998; Zanetti et al., 2001; van Halteren-van Tilborg et al., 2007; Kessels et al., 2011; De Wit et al., 2021, 2022).

Motor learning has traditionally been regarded as a form of non-declarative memory. It is defined as the process of (re)gaining or retaining a given level of motor performance (Krakauer et al., 2019). Indeed, AD patients are still able to (re)learn motor tasks, although learning success depends on the type of task (Willingham et al., 1997; van Tilborg and Hulstijn, 2010), how feedback is provided (van Halteren-van Tilborg et al., 2007), whether rewards are present or not (Wong et al., 2019) and how engaged the patient is in the task (Laffan et al., 2010).

More recently, it has been suggested that motor learning not only relies on non-declarative memory but also involves declarative processing (Mazzoni and Krakauer, 2006; Keisler and Shadmehr, 2010; Taylor and Ivry, 2011; Taylor et al., 2014; McDougle et al., 2022) The arguments find their basis in a computational theory of error-based motor learning. This theory proposes that the learning process involved in acquiring a new mapping between motor commands and behavioral outcome involves a fast adaptive process that learns quickly but also decays rapidly and a slow process that learns slowly but has good retention (Smith et al., 2006). Behavioral and neural evidence for such dual-rate learning has been reported for both force field and visuomotor adaptation using spontaneous recovery paradigms (Lee and Schweighofer, 2009; Trewartha et al., 2014; Inoue et al., 2015; Sarwary et al., 2018).

Because the fast process is mainly driven by large movement errors, Keisler and Shadmehr (2010) argued that it shares resources with the declarative memory process. They reasoned that large movement errors enter the learner’s awareness and are thus explicitly processed (Malfait and Ostry, 2004), and hence engage the declarative memory system. In support, the authors showed that performing a declarative memory task (word-list recall) after completion of a reach adaptation task produced interference with the memory of the fast process, but not the slow process (Keisler and Shadmehr, 2010). Taylor et al. (2014) showed that implicit adaptation demonstrated slow adaptation dynamics, while explicit adaptation demonstrated fast adaptation dynamics. Finally, it has been shown that limiting movement preparation time suppresses the recruitment of explicit processing, such that learning is best described by a single implicit process (Fernandez-Ruiz et al., 2011; Haith et al., 2015; Leow et al., 2017).

Therefore, if explicit, declarative processing affects the ability of the fast process to lay down motor memories, it can be hypothesized that its retention rate is lower in AD patients than in controls. Here, after careful neuropsychological examination, we tested 21 early-stage AD patients in a spontaneous recovery paradigm using a force field adaptation task with reaching movements. Compared to age and education level matched healthy control participants, their data show that the fast, but not the slow adaptive process is affected, which not only refines computational theories of motor learning but also allows for possible clinical translation.

## Materials and Methods

### Participants

The medical-ethics committee of the Radboud University Medical Center judged this study exempt from formal medical-ethical approval according to the WMO Act (CMO Arnhem-Nijmegen 2017-3162) due to the minimal burden it imposed on participants. The present study was subsequently approved by the ethics committee of the Social Sciences faculty of Radboud University in Nijmegen, The Netherlands (ECG2017-0805-504). All participants (patients and controls) gave written consent to participate in the study and were reimbursed for their time at a rate of 10€ per hour.

All participants had normal or corrected-to-normal vision. The Edinburgh Handedness Inventory (Oldfield, 1971) showed that all but one participant (from the control group) were right-handed. Participants indicated their education level based on the Dutch educational system (range 1–7, with 1 = less than primary school; 7 = academic degree; (Verhage, 1964)).

Patients were recruited from the memory clinic of the Radboud University Medical Center in the period between May 2017 and May 2019. Inclusion criterion for the patients was: having a declarative memory impairment verified by neuropsychological assessment due to Alzheimer’s disease (either amnestic Mild Cognitive Impairment [aMCI] or mild dementia). The exclusion criteria were a history of other neurological diseases that affect the brain (stroke, Parkinson’s disease, brain tumor), a history of or an active psychiatric disorder (including psychotic disorders or a substance-use disorders), and no command of Dutch language. The main experimental group consisted of 20 patients (5 women, aged 60-87) who were all diagnosed with (amnestic) MCI due to Alzheimer’s disease or a mild Alzheimer’s dementia. MCI due to Alzheimer’s disease refers to the symptomatic predementia phase of AD. In MCI patients, the degree of cognitive impairment is not age-appropriate (Smith et al., 1996; Albert et al., 2011). Clinical diagnoses were established based on a multidisciplinary assessment at the memory clinic of the Radboud University Medical Center, and were supported by a clinical interview with the patients and their informants, neuroradiological findings, neuropsychological assessment and a review of the patients’ medical history, in accordance with current criteria (Albert et al., 2011; McKhann et al., 2011). The clinical dementia rating (CDR, Hughes et al. 1982) of the patients was 0.5 (MCI) or 1 (mild dementia).

As a control group we recruited 25 age- and education level matched, cognitively unimpaired participants (Verhage, 1964). Four participants from this group were excluded from the analysis: one due to failure to follow task instructions, one due to experienced pain in the right arm when holding the robot handle and two due to failure in experimental recording; hence 21 healthy participants (13 women, aged 61-87) were included in the analysis. This group was both age-matched (*t*(39) = 0.5, *p* = 0.6, t-test) and matched at the education level (*t*(30.31) = 1.2, *p* = 0.26, t-test). Visual inspection of the data of the left-handed participant, who performed the task with the right hand, did not reveal differences from right-handed participants’ data and therefore was included in the analysis. The demographic (and neuropsychological) characteristics of the AD patients and the control group are presented in Table 1.

**Table 1.** Demographic and neuropsychological characteristics of the AD patients and the control participants. Data are reported as mean (SD) or number. CDR – Clinical Dementia Rating, 0.5-1 indicates very mild to mild dementia. MoCA – Montreal Cognitive Assessment, maximum score 30, education adjusted (1 extra point for those with 12 or less years of education). MoCA-MIS – Montreal Cognitive Assessment Memory Index Score, maximum score 15. NART – National Adult Reading Test, intelligence quotient estimate, maximum score 130. Doors test, visual recognition memory task, maximum score 24. Education level: 1 = less than six years elementary school; 2 = six years elementary school; 3 = more than six years elementary school; 4 = vocational training; 5 = community college; 6 = advanced vocational training; 7 = university degree.

|  | **AD patients** | **Control participants** |
| --- | --- | --- |
| **Age (years)** | 75.6 (6.5) | 74.4 (7.4) |
| **Number of men/women** | 15/5 | 9/12 |
| **Education level (1-7)** | 5.0 (1.9) | 5.6 (1.1) |
| **Education in years** | 13.7 (5.7) | 15.6 (3.7) |
| **Number of patients with CDR 0.5/ 1** | 6/14 | n/a |
| **MoCA Total score (points)** | 20.1 (3.9) | 26.4 (1.9) |
| **MoCA-MIS (points)** | 3.5 (3.5) | 12.7 (1.9) |
| **NART (IQ)** | 101.2 (15.0) | 108.6 (7.4) |
| **Doors test (points)** | 12.0 (3.9) | 17.9 (3.2) |

### Neuropsychological testing

All participants performed a set of neuropsychological tests before the reaching task. First, participants performed the Dutch version of the National Adult Reading Test (NART) to estimate their premorbid level of intellectual functioning (Schmand et al., 1992). Second, the Montreal Cognitive Assessment (MoCA) with the cued-recall and multiple-choice memory test items (enabling calculating the Memory Index Score, MoCA-MIS) was administered to assess global cognitive functioning and memory (Nasreddine et al., 2005; Julayanont et al., 2014). Finally, participants completed the Doors Test (parts A and B), a visual recognition memory test from the Doors and People Test (Baddeley et al., 1994) to assess declarative memory. The results of the neuropsychological characteristics of the patients and control participants groups are presented in Table 1.

An independent samples *t*-test on the IQ estimates (based on the NART) did not reveal any significant differences between the two groups (t(27.5) = 2.0, p = 0.057, t-test). However, as expected, the AD patients had worse global cognitive functioning than the control group (MoCA: t(27.2) = 6.6, p < 0.001, t-test) and worse declarative memory performance (Doors Test: t(39) = 5.3, p < 0.001, MoCA-MIS: t(28.4) = −10.3, p<0.001, t-test).

### Experimental Setup

During the experiment, participants sat in front of a planar robotic manipulandum (Howard et al., 2009). While holding the handle of the manipulandum, they performed reaching movements in the horizontal plane with their right arm. Participants did not have direct visual feedback of the arm due to a semi-silvered mirror that covered the arm (Figure 1A). An air sled underneath the right forearm allowed frictionless movements. All visual stimuli were presented on an LCD monitor (model VG278H, Asus) that was viewed via the semi-silvered mirror. The refresh rate of the display was 120 Hz. Stimuli were shown in the same plane as the movements. Hand position, derived from the robot’s handle position, was presented as a cursor (red disc, 0.35 cm radius). The target (yellow disc, 1 cm radius) was placed 12 cm out in the straight-ahead direction from the central home position (white disc, radius 1 cm). The home position was located about 30 cm from the participant’s chest. An auditory imperative stimulus and warning/error message were presented via speakers that were behind the workspace of the experiment. Robot handle position data were measured at 1000 Hz.

**Figure 1.**
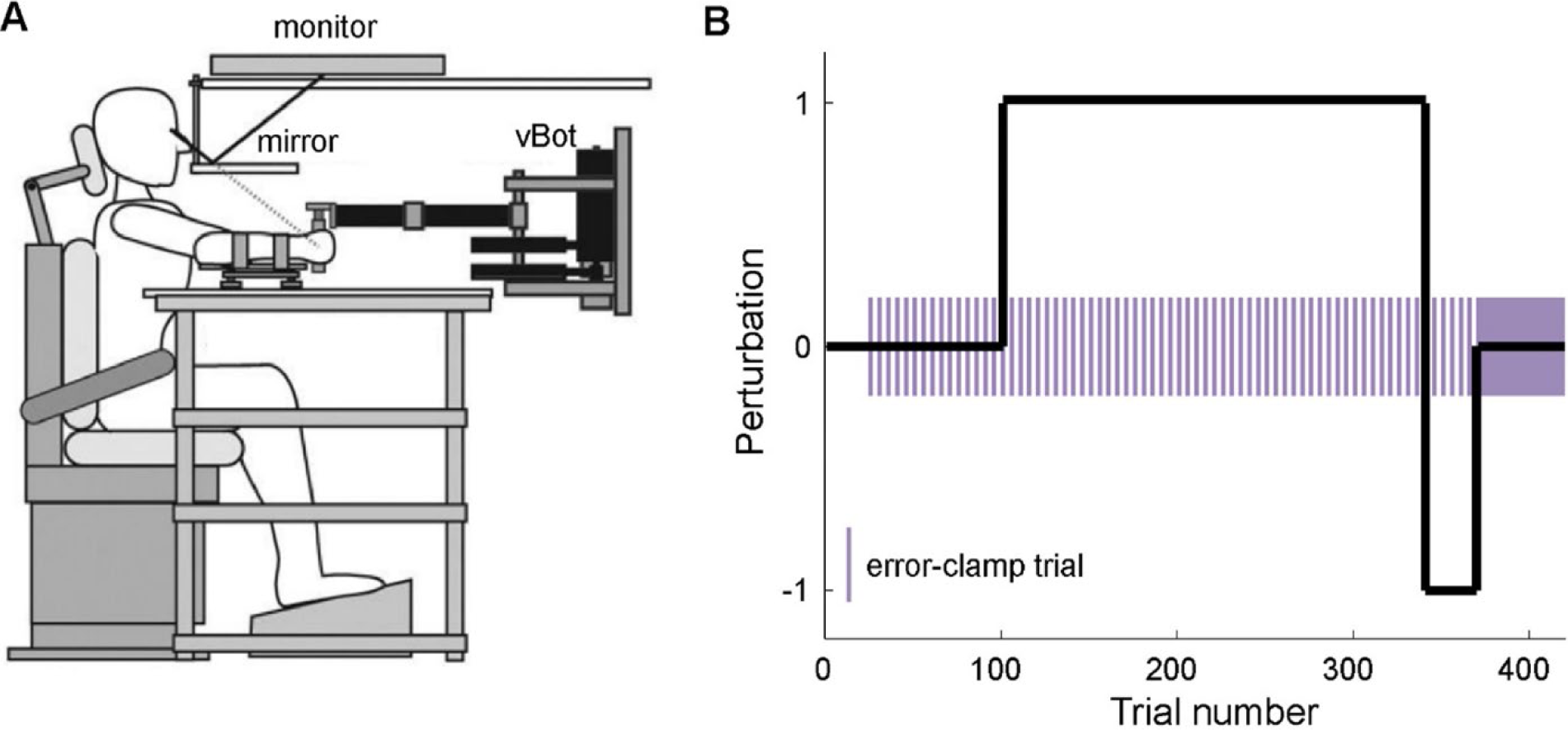
A. Experimental setup. Participants held the handle of a planar robotic manipulandum (vBot) with their right hand while their arm was resting on an air sled floating on a glass-top table. Visual stimuli were presented through a mirror. Image reproduced from Franklin and Wolpert (2008) with permission. **B.** Experimental paradigm. The experiment started with 100 null-field trials; these were followed by 240 clockwise force field-trials, 30 CCW force-field trials and finally 50 error clamp trials. To track the progression of learning, every 5^th^ trial from trial 21 to 370 was an error clamp trial (purple bars).

There were 3 types of trials: 1) null-field trials (robot forces were turned off); 2) curl force field trials and 3) error clamp (EC) trials. In curl force field trials, the robot produced forces that were perpendicular to the movement direction and proportional to the reach speed:

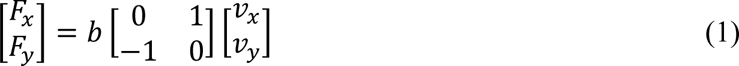

where *x* and *y* are the lateral and sagittal directions, *F*_*x*_ and *F*_*y*_ are the robot forces applied at the hand, *v*_*x*_ and *v*_*y*_ are hand velocities, and *b* is the field constant (± 13 Ns/m). The sign of the field constant determined the direction of the force field. Error clamps served to measure the adaptation index (AI). In error clamp trials, the hand was constrained to a straight path from the start to the target with a spring constant of 6,000 N/m and a damping constant of 7.5 Ns/m. At the end of each reach, participants were asked to relax their arm while the arm was passively returned to the start position following a minimum jerk profile with a duration of 700 ms.

### Experimental Paradigm

The reaching task was first explained to the participant by the experimenter using a paper version of the task workspace and a dummy robot handle at the same desk where the neuropsychological tests were administered. Once the participant understood the task, they were asked to take a seat at the desk where the reaching experiment took place. To start a trial, participants placed the cursor inside the home disc. After 500 ms, the yellow target appeared, which was accompanied by an auditory beep. Participants were instructed to move in a straight line to the target as soon as it appeared. The target turned green if the cursor was in the target. At the end of the reach, the robot returned the hand to the start position. Inter-trial-interval was set at 200 ms. In order to get accustomed with the experimental setup, participants performed 10 practice trials with the vBot before the experiment. If necessary, this practice block was repeated.

The experiment started with 100 null-field trials (Figure 1B). Thereafter the clockwise curl force field was turned on for 240 trials. This was followed by 30 trials of counter-clockwise curl force-field and finally participants performed 50 error clamp trials. From trial 21 to 370 the null-field and force-field trials were interleaved with error clamp trials (every 5^th^ trial).

To ensure that participants did not slow down the reaching pace along the experiment, a warning message “Move faster” was given at the end of the reach if movement duration (time between movement onset and reach offset) was longer than 500 ms. A warning message “Stay in the target!” was displayed if the cursor left the 3 cm radius area around the target disc within 200 ms after entering it. Warning messages did not lead to re-starting or exclusion of the respective trial. Participants received an error message if they did not start the reaching movement within 1000 ms and the trial was re-started. All warning and error messages were displayed for 1250 ms.

### Data analysis

Analyses were performed using Matlab 2018b (MathWorks). Reach onset was determined as the time point when the cursor speed exceeded 5 cm/s. Reach offset was determined as the time point 200 ms after the distance between the cursor and the center of the target disc was smaller than 3 cm. There were two exceptions to this criterion: 1) If the cursor after entering the 3 cm radius area around the target disc center left it again within 200 ms. In such cases, reach offset was determined upon exiting the 3 cm radius area; 2) If the cursor speed dropped below 5 cm/s outside the 3 cm radius area from the target, reach offset was determined at this timepoint.

Trials in which the participant moved less than 6 cm from the middle of the starting position were removed from further analysis. For each error clamp trial, we computed the adaptation index (AI), which represents the fraction of ideal force compensation in response to the curl force-field. To this end, based on the velocity of the handle along the channel, we calculated the time-varying lateral force that would have been generated by the force-field in a field trial. The force measured in the error clamp was regressed against this theoretical force, providing the regression coefficient, which was taken as the adaptation index (see Joiner and Smith, 2008 for more details). Adaptation indices during the force-field trials were baseline corrected by subtracting the mean AI during the null trials in the beginning of the experiment.

### State-space modeling

We fitted a Bayesian hierarchical version of the dual-rate model (Smith et al., 2006) to the time course of the AI across the various phases of the experiment. According to this model, the total adaptation at trial *n* + 1 can be represented by the sum of the states of a fast process (*x*_*f*_ (*n* + 1)) and a slow process (*x*_*s*_(*n* + 1)),

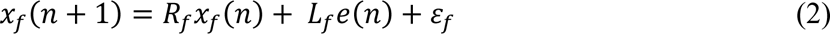

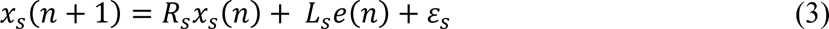

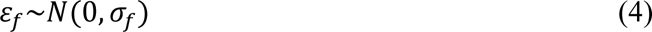

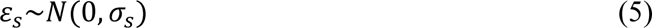

The state of each process depends on the state at the previous trial *n*, multiplied by a retention rate (*R*), and the error of the previous trial, *e*(*n*), multiplied by the learning rate (*L*), while also Gaussian state noise (*ε*_*f*_, *ε*_*s*_, Eq 4 and 5) is added at each trial. Both processes have independent retention and learning rates, which were constrained as follows: 0 < *R*_*f*_ < *R*_*s*_ < 1 and 0 < *L*_*s*_ < *L*_*f*_ < 1.

The sum of the fast and the slow process is the net motor output (*x*(*n* + 1)),

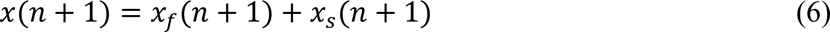

The actual movement output (*y*(*n* + 1)) is equal to the sum of the net motor output (Eq 6) and Gaussian output noise (*ε_output_*),

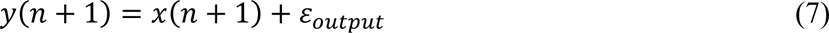

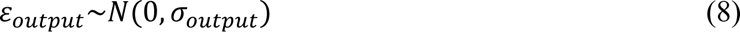

The error (*e*(*n*)) on a trial (*n*) is defined as the difference between the actual movement output, *y*(*n*), and the applied perturbation force, *f*(*n*),

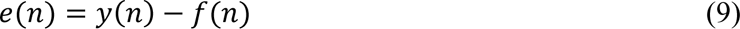

We used a hierarchical implementation of this dual-rate model, following Ferrea et al. (2022), to estimate the learning and retention rates of individual participants. The advantage of using hierarchical modeling is that the estimate of the individual parameters is informed by data from all other individuals in the same group (patients vs. controls) (Kruschke, 2015).

Figure 2 shows a graphic diagram of the full hierarchical model. Each participant’s learning and retention rate for both the slow and fast process were sampled from priors: normal distributions truncated to the interval [0, 1]. The means *μ* and standard deviations *σ* of these priors were hyperparameters of the hierarchical model, which were sampled from hyperpriors. For the means *μμ*, the hyperpriors were normal distributions truncated to the interval [0, 1]. The means of these hyperpriors were 0.998, 0.85, 0.1 and 0.1 for *R*_*s*_, *R*_*f*_, *L*_*s*_, and *L*_*f*_, respectively, and the standard deviations were 0.01, 0.5, 0.5 and 0.5 for these respective parameters. For the standard deviations of the priors, the hyperparameters were half-Cauchy distributions. For all learning and retention rates, these were Cauchy distributions with location parameter 0 and scale parameter 0.5, truncated to values > 0. The standard deviation *σ* of both the fast and the slow state noise and the standard deviation of the output noise were constrained to values > 0 and these were sampled from normal priors with hyperparameters mean *μ* and standard deviation *σ*. All these hyperparameters were sampled from half-Cauchy hyperpriors. For the means, the location and scale parameter were 0 and 1, respectively, whereas these were 0 and 0.5 for the standard deviations.

**Figure 2.**
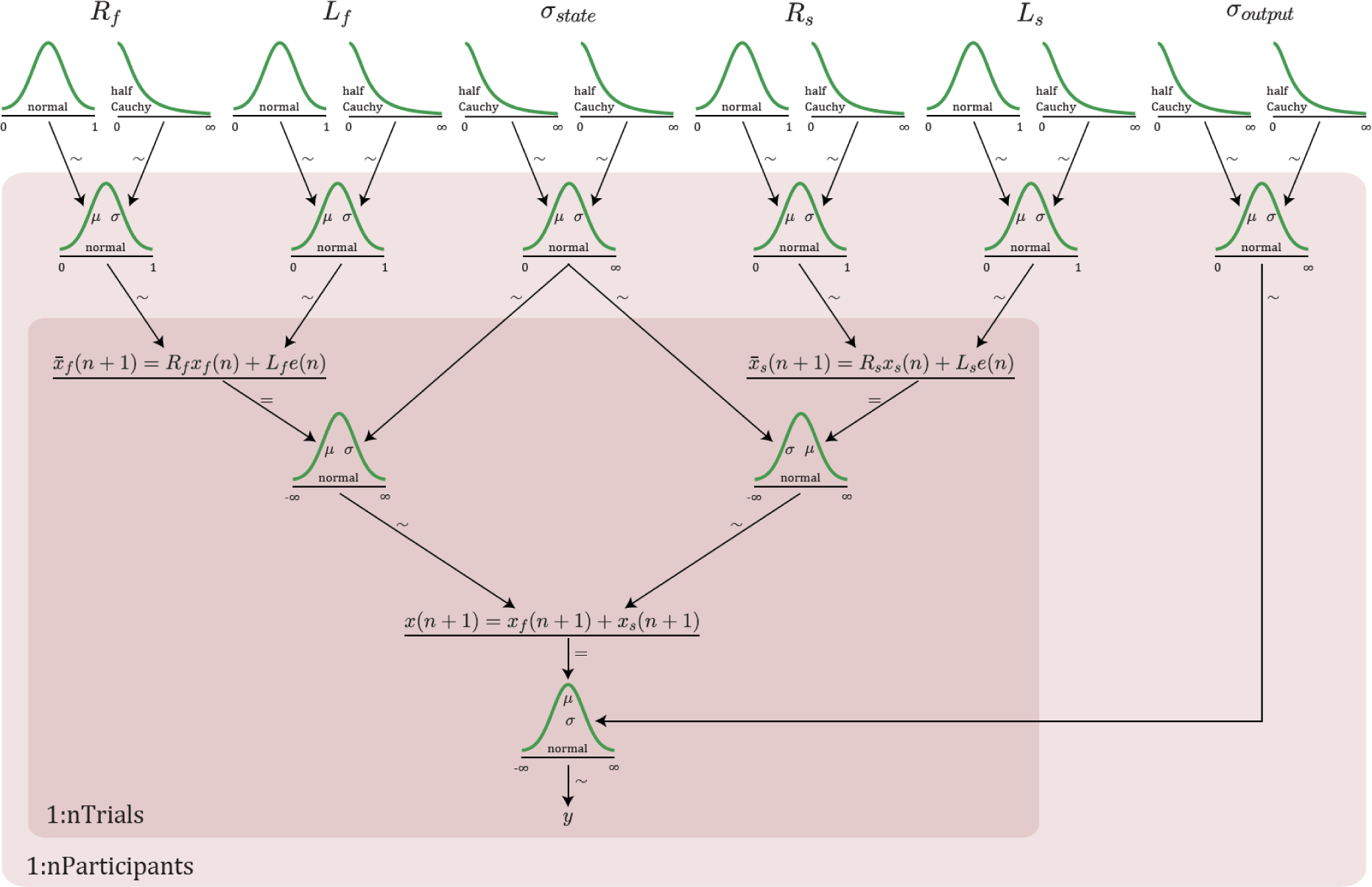
Diagram of the hierarchical implementation of the dual-rate model. Individual participant’s learning and retention rate estimates for both the slow and fast process were sampled from truncated normal distributions (the priors shown at the second row), which were defined in terms of their mean *μ* and standard deviation *σ*. These parameters were hyperparameters that were sampled from the hyperpriors in the top row (truncated normal distributions for the means, and half-Cauchy distributions for the standard deviations). The standard deviations *σ* of the fast and the slow state noise and the output noise were sampled from normal priors (in the second row). Their mean *μ* and standard deviation *σ* were sampled from half-Cauchy hyperpriors in the top row. The distributions on the white background were sampled at the group level; those on light red background at the participant level and those on the dark red background at the trial level.

To have the parameter estimates mostly determined by the data, all hyperpriors were fairly uninformative, except for those pertaining to the means of the retention rates, which were chosen more informative to speed up the convergence of the Markov chains. We verified that making these hyperpriors less informative also led to the same parameter estimates, but at the expense of a much lower effective sample size.

We used independent hierarchical models for the AD patients and for the healthy controls. Importantly, the same model with the same priors was used for both groups.

### Statistical analysis

Statistical analyses of the outcomes of the neuropsychological tests and the adaptation process with AIs were performed using SPSS Statistics 25. To compare the adaptation process along the reaching experiment we compared the AIs of the AD patients and control participants in three different phases of the experiment: 1) at the plateau of adaptation during the clockwise force field (last 15 EC trials); 2) during counter clockwise force field (last 3 EC trials); 3) and finally, at the beginning of the final EC phase at the end of the experiment (first 12 EC trials). To this aim we first calculated the mean AI for each phase per participant and compared groups with paired samples *t*-tests.

Parameter estimation of the dual-rate model was conducted using Markov Chain Monte Carlo (MCMC) methods in Stan (CmdStan version 2.17.1) via its Matlab interface MatlabStan. The MCMC method gives representative samples of the posterior distribution of the model parameters given the data. We ran the model on 4 chains with a burn-in phase of 3,000 samples and 25,000 iterations for each chain. No thinning was used. We inspected the following MCMC diagnostics for each parameter: the convergence diagnostic 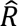, the effective sample size (ESS) and the Monte Carlo Standard Error (MCSE). For all hyperparameters, the fits revealed the 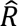 < 1.022, the ESS > 250 and the MCSE < 0.0023. Supplementary Table 1 lists the 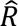, ESS and MCSE for all hyperparameters.

We performed a posterior predictive check to assess whether the model gave valid predictions of the data. A grand-average visualization (Figure 4) was generated from averaging the posterior predictive checks for all participants, separately for patients and healthy controls. Individual posterior predictions were generated from random draws (*n* = 500) of the posterior distributions of the estimated learning and retention rates of the fast and the slow process, while the state and output noise was drawn from the corresponding normal distributions.

To assess whether there were any group differences for each of the main four model parameters, we performed Region of Practical Equivalence (ROPE) analysis using the respective hyperparameters. To this end, we determined for all retention and learning rates the posterior distribution of the difference between the hyperparameter reflecting the mean of the AD group and that of the healthy control group. We tested the ROPE for each parameter separately using the formula (Kruschke, 2018):

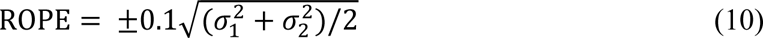

in which 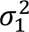 and 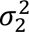 are the variances of the posterior distributions of the two groups. We accepted that there was no difference between groups if the 95% highest density interval (HDI) of the posterior distribution of the difference between groups fell completely within the ROPE. We rejected that there was no difference between groups if 95% HDI fell completely outside the ROPE. In all other cases, we withheld a decision.

### Data availability

Upon publication, all data and code will be available from the data repository of the Donders Institute for Brain, Cognition and Behaviour via the following URL <TO be added>.

## Results

Patients and controls made forward reaching movements to a visual target, using a protocol consisting of four blocks. A baseline phase of 100 trials was followed by a long phase of 240 trials during which they were exposed to a clockwise (CW) force field (except for the intervening error clamps). This was followed by a shorter exposure of 30 trials to a counterclockwise (CCW) force field before the final error clamp block of 50 trials began.

### Force-field adaptation and spontaneous recovery in AD

To quantify adaptation within the first three blocks, every fifth trial was an error clamp trial (from trial 20 in the first baseline block), in which the robot clamped the reach to a straight line, while the compensatory force was measured, from which the adaptation index was computed (see Materials and Methods). All trials in the fourth block were error clamps.

All patients and control participants who were included in the study were able to perform the task in accordance with the test instructions. Figure 3 shows the adaptation index for AD patients and healthy controls during the four experimental blocks. As expected, force expression was unsystematic during the baseline phase since the robot did not perturb the reaches. In both groups, the AI gradually increased during exposure to the CW force field suggesting that like controls, patients also learned to compensate for the forces, and approached an asymptotic level. While the asymptote was below one in both groups, indicating that neither patients nor controls completely cancelled the force applied by the robot, the patients compensated slightly but significantly less (*t*(39) = −2.06, *p* = 0.046, t-test). During the subsequent CCW block, when the force field had switched to the opposite direction, the AI quickly returned to baseline levels and for the controls even switched sign while starting to compensate for the CCW force field. This indicates that patients did not adapt as much to the second force-field as the control participants (*t*(39) = 2.97, *p* = 0.005, t-test). Next, during the final error clamp block, the AI rapidly rebounded, in both patients and controls, expressing part of the compensatory strategy for the first force field, known as spontaneous recovery. The AI was initially larger in patients than controls (*t*(39) = 3.91, *p* < 0.0005, t-test), but both groups plateaued at about the same level during the end of the error clamp phase. We next adopted a modeling approach to interpret this altered pattern of spontaneous recovery.

**Figure 3.**
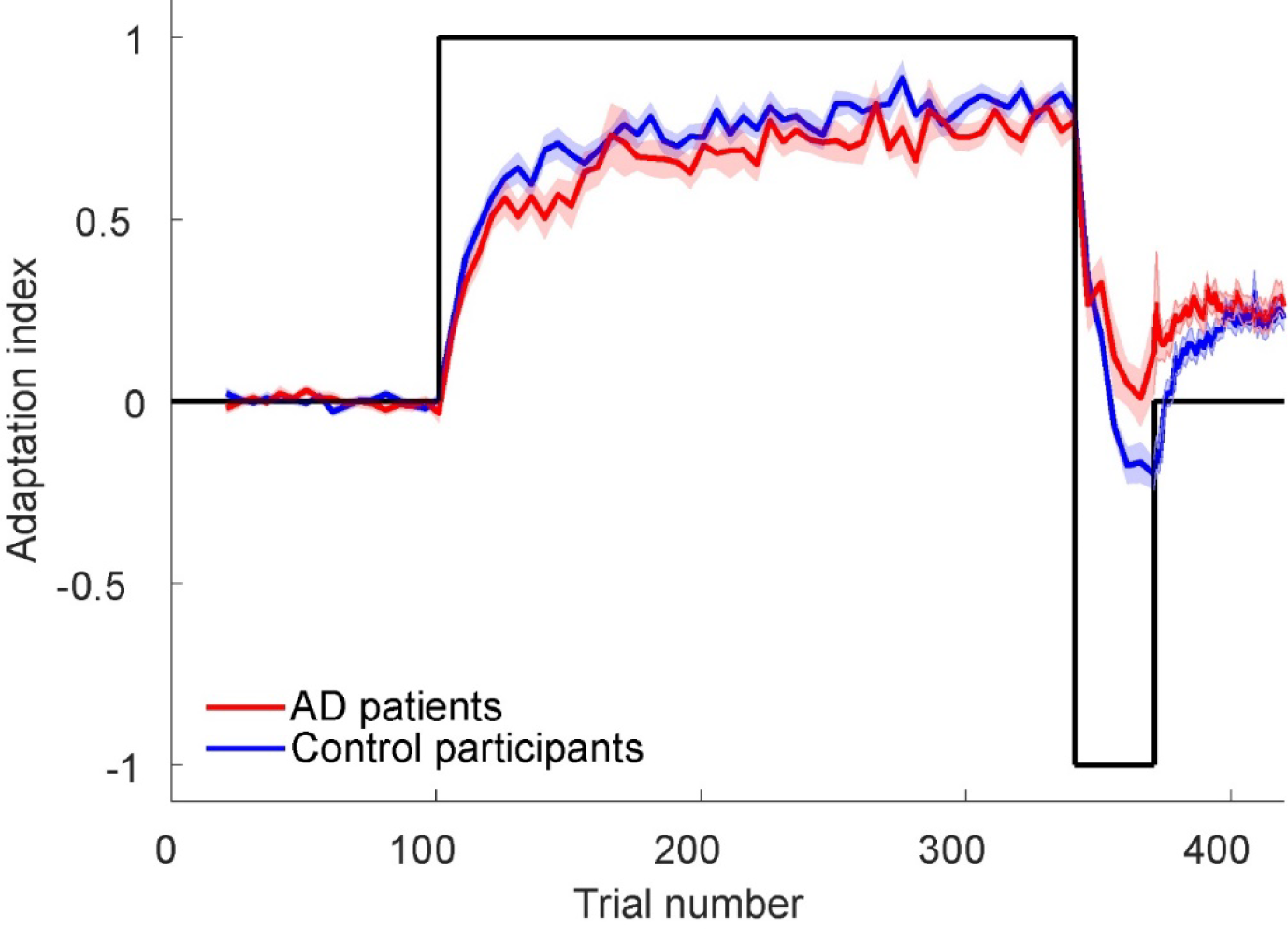
The average adaptation index as a function of trial number for control participants (blue) and AD patients (red). The black line indicates the direction of the force field (CW/CCW). The shaded area denotes ±SE.

### The fast, not the slow adaptive process, is affected in AD

We fitted a Bayesian hierarchical version of a dual-rate adaptation model (Smith et al., 2006) to capture the time course of the AI. Figure 4 illustrates the two internal states of the model. The state of the fast process (in green) demonstrates quick learning and quick decay; the slow process (in light blue) illustrates slower learning and hardly decay. The model accounts for the main characteristics of the measured AI time courses, in both patients and controls. Upon closer inspection, however, there are subtle differences. First, both patients and controls generally adapt faster than the model implies. Second, the model does not fully capture the final level of spontaneous recovery. For both groups, the model shows lower spontaneous recovery than the data. Despite these subtle differences between data and model fits, the differences between the groups in the data are also visible in the posterior predictive checks of the model.

**Figure 4.**
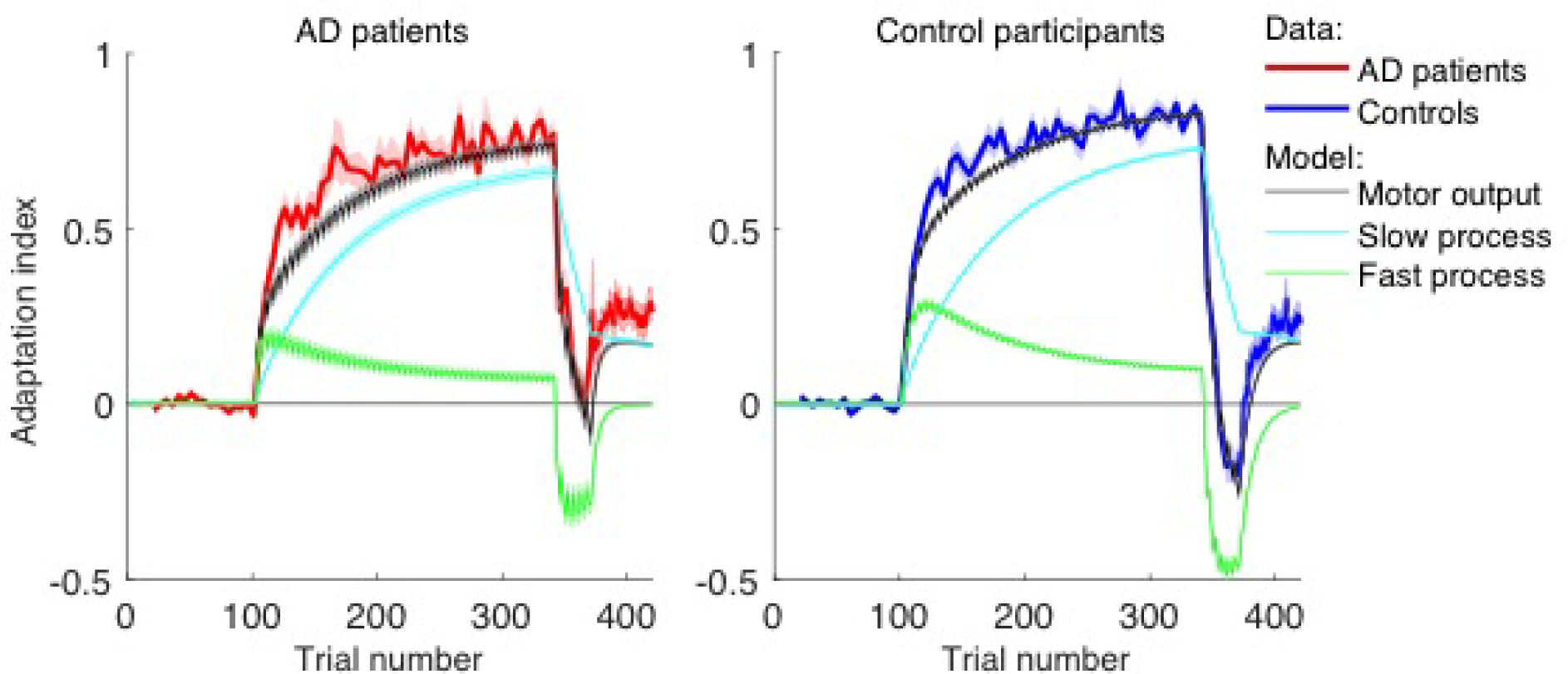
Model fit. Adaptation index as a function of trial number for the patients with Alzheimer’s disease (red; left panel) and control participants (blue; right panel). The fitted motor output (black) as a result of the slow process (cyan) and the fast process (green) is the mean of the posterior predictive check performed for individual subjects (500 random draws each). The shaded area denotes ±SE. The zigzag pattern in the model predictions during the adaptation phases arises from the succession of force-field trials (series of 4 trials during which adaptation increased) and error-clamp trials (single trials during which adaptation decayed). The data do not show the zigzag pattern because they show only the adaptation index during the error-clamp trials (the only trials in which the adaptation index was measured).

Figure 5 illustrates point estimates of the fitted parameters (based on the mean of the posterior distributions) of the hierarchical implementation of the two-state model for individual patients and controls, while the mean and the full posterior density of the hyperparameters representing the population mean of each parameter are also shown. We used a ROPE analysis to test whether there were differences in these hyperparameters between the groups. Consistent with our hypothesis, *R*_*f*_ was the only parameter for which we can reject the null value: AD patients had a lower retention rate of the fast process than control participants (Figure 5). For the other parameters (*R*_*s*_, *L*_*s*_, and *L*_*f*_), the ROPE fell completely inside the HDI of the posterior distributions of the parameters, indicating that the decision about these parameters is withheld. For the parameters of the slow process, this is partly due to the relatively broad densities of the hyperparameters. These results suggest that the differences in motor adaptation between the AD patients and controls can be explained by a reduced retention rate of the fast process in AD patients. Supplementary Table 2 lists the means and the HDIs for all parameters.

**Figure 5.**
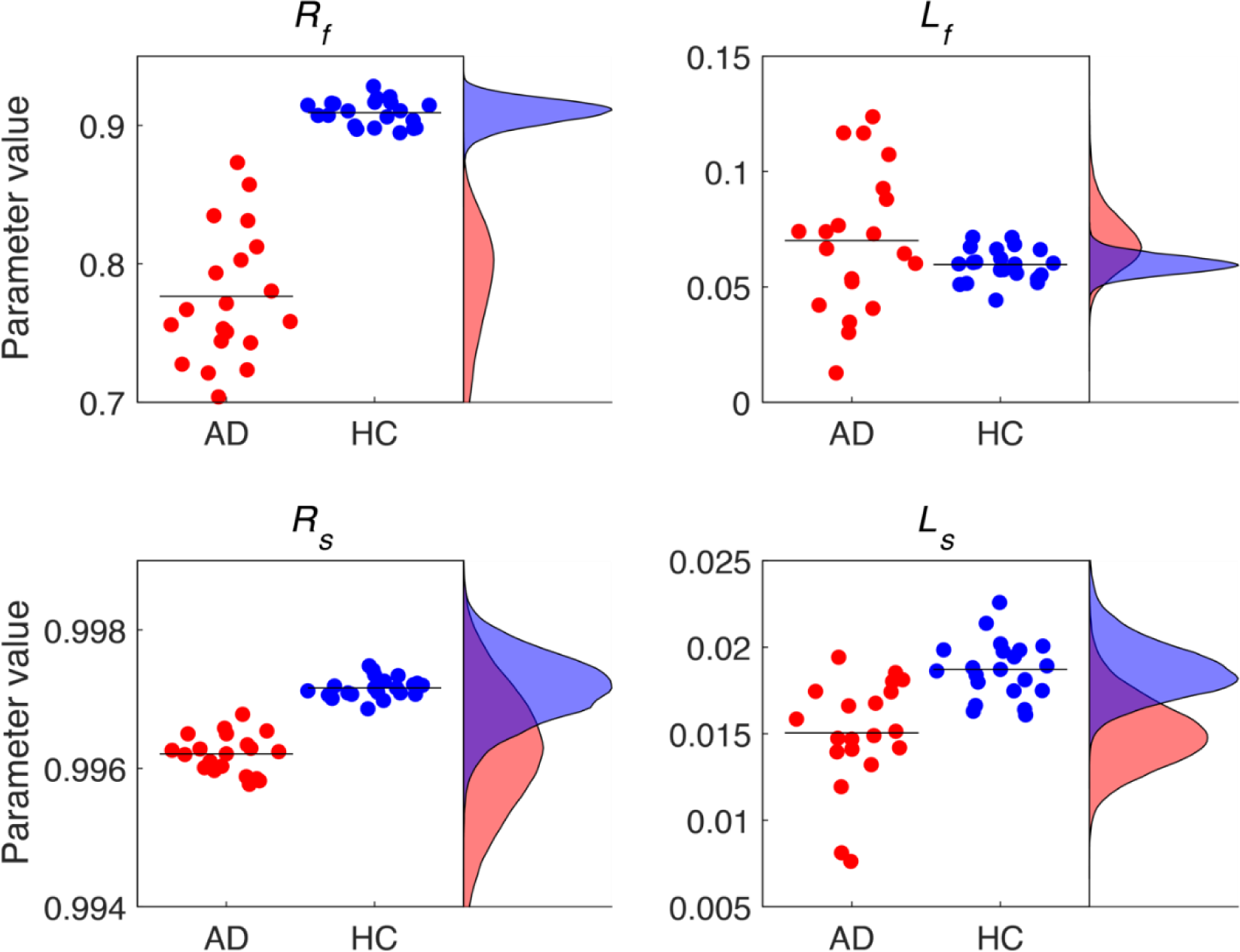
Individual parameter estimates (dots) and mean (black horizontal lines) and full posterior densities (on the right hand side of each plot) of the hyperparameters representing the population mean of each parameter for AD patients (AD, red) and healthy control participants (HC, blue).

The conclusion that only the slow retention rate differed between the groups can fully explain the differences in the observed adaptation curves. Figure 6 shows simulations of the dual-rate model for two sets of parameters, representing AD patients and control participants. The parameters that did not differ between the groups were for both simulations set to the average estimate of both groups (*L*_*s*_ = 0.0169, *L*_*f*_ = 0.0649, *R*_*s*_ = 0.9967). The retention rate of the fast process was set to the mean of each experimental group (*R*_*f*_ = 0.7765 for the AD patients, *R*_*f*_ = 0.9092 for the controls). The simulated controls adapt more rapidly to the first force field. This is because the fast process, which drives the initial adaptation, has better retention for this group than for the simulated patients. Another consequence is that the adaptation level of the fast process is lower for the simulated patients than for the controls, which in turn leads to *stronger* adaptation of the slow process for the simulated patients, and almost the same net adaptation for both groups at the end of the adaptation phase. Due to their better retention, the fast process of the simulated controls adapted more strongly to the second force field than that of the simulated patients. Since the adaptation level of the slow process did not differ much between the groups, the net adaptation at the end of this phase was strongly negative for the simulated controls and close to zero for the simulated patients, consistent with the data. Finally, both simulated groups reached the same final level of spontaneous recovery, but the simulated patients reached this level earlier than the simulated controls, also consistent with the data.

**Figure 6.**
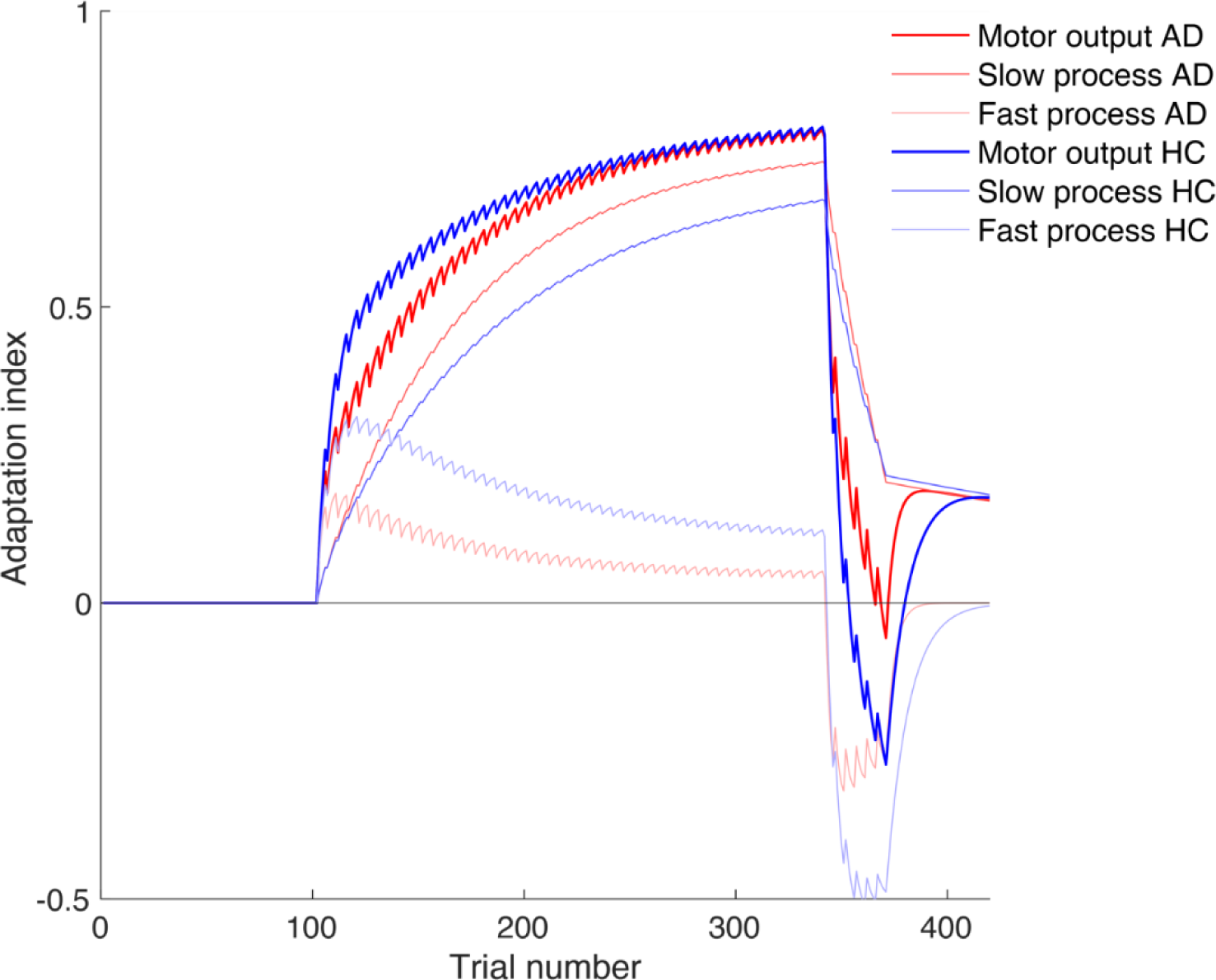
Simulation of the dual rate model for two sets of parameters, representing AD patients and healthy control (HC) participants. For both groups: ***L_s_* = 0.0169, *L_f_* = 0.0649, *R_s_* = 0.9967, AD patients: *R_f_* = 0.7765, HC: *R_f_* = 0.9092**. All noise levels were set to zero.

## Discussion

We measured the time course of reach adaptation of early-stage AD patients while they were exposed to perturbing force fields in the context of a spontaneous recovery paradigm. Guided by a dual-rate adaptation model, we reverse engineered the mechanisms underlying their motor learning ability in terms of a slow and a fast adaptive process.

Our results show that patients with declarative memory impairment compensated slightly less than healthy controls for the force perturbations, although both groups showed clear signs of spontaneous recovery. As Smith and colleagues demonstrated, effects of spontaneous recovery can be explained by the concurrent existence of multiple adaptive processes (Smith et al., 2006). This is confirmed by our dual-rate model analysis, which further shows that the fast, but not the slow process is affected in AD. More specifically, the retention rate of this process – the recall of this state from trial to trial - was significantly lower than in controls, while the learning rate of the process – the error sensitivity – did not differ significantly.

In the motor learning literature, there is substantial behavioral evidence that explicit processes can contribute to motor adaptation (Heuer and Hegele, 2008; Anguera et al., 2010; Krakauer et al., 2019). Recent neuroimaging work corroborates these observations by demonstrating bidirectional interactions between neural substrates of motor memory and declarative memory (Kim, 2020), which change with age (Wolpe et al., 2020). The present findings constrain the role of the declarative memory to interactions with the fast adaptive process. This inference here is based on a clinical population with declarative memory impairment but concurs with findings in healthy participants using other paradigms. For instance, Keisler and Shadmehr (2010) demonstrated that a word-pair learning task, which taxes declarative memory, interferes with the memory of the fast, but not slow process. Taylor et al. (2014) showed that explicitly-declared performance reports during motor learning follow fast adaptation dynamics.

From a clinical perspective, it is commonly thought that patients with declarative memory deficits can learn some motor tasks, such as rotary pursuit, serial reaction time tasks and mirror-tracking (Ferraro et al., 1993; Willingham et al., 1997; Dick et al., 2001; Rouleau et al., 2002; Hong et al., 2020; De Wit et al., 2022), because such tasks rely on implicit ways of learning. For example, Shadmehr et al. (1998) showed that an amnesic patient (H.M.), due to a resection of the medial-temporal lobe, was still able to adapt his reaches to a perturbing force field, but was unaware that he learned the task. Noteworthy, the adaptation in this patient proceeded very slowly, as if only a slow process was involved and the fast adaptive process was lacking. Our memory-impaired patients still show evidence for the operation of a fast process, suggesting that the explicit processes are only partially affected. This may not be surprising given their mild declarative memory impairment, as indicated by lower MoCA-MIS and Doors Test scores in AD patients compared to the controls. Whether the contribution of the fast process would be smaller in AD patients at a more advanced disease stage remains to be investigated.

Recently, McDougle et al. (2022) described another case report of a patient (L.S.J.) with near-complete bilateral loss of her hippocampi. She was tested on various reach adaptation tasks, including a force-field paradigm for evoking spontaneous recovery. L.S.J. showed robust learning, but weaker than controls, as well as spontaneous recovery, all consistent with the present observations. Furthermore, dual-rate modeling showed that L.S.J. differed from controls in the retention parameter (*R*_*f*_) of the fast motor learning process. The authors found this parameter to be increased, which they explain by increased rigidity of the fast process, reflecting reduced flexibility. In contrast, all our 20 AD patients showed a lower retention rate of the fast process compared to the controls, which can be expected assuming this parameter characterizes their amnesia.

We can only speculate about this difference with the study by McDougle and colleagues. Experimentally there seem to be only small differences (e.g. in the number and proportion of error clamp trials), but we tested a selective patient group (AD patients with mild cognitive impairment) which may be different than their individual case due to the neurodegenerative nature of AD. Furthermore, we used a hierarchical Bayesian modeling procedure to fit the dual rate model, which is an advanced approach that deviates from McDougle et al.’s and other previous approaches (Albert and Shadmehr, 2018; Coltman et al., 2021). In this approach the estimate of each of the four parameters (*R*_*s*_, *R*_*f*_, *L*_*s*_, and *L*_*f*_) of each individual is simultaneously informed by the data from the other individuals, because all individuals inform higher-level parameters, called hyperparameters, which constrain all the individual parameters making them less sensitive to individual outliers (Kruschke, 2015). In other words, the hyperparameters constrain the estimates of the individual parameters of the retention and learning rates.

Based on an HDI+ROPE decision rule, the fast retention rate marked a significant neurocomputational alteration at the group level as well as at the level of each individual patient. For none of the other three parameters this was found to be the case. Since a lower fast retention rate was robustly found in each patient, it opens up the possibility to use it as a prognostic or diagnostic marker to complement traditional taxonomies or neuropsychological test or to use it to track the development of the disease.

A further point of discussion is that only the retention rate of the fast process was changed in our patients, not the learning rate. This suggests that sensitivity to error of the fast process is not changed in our patients, but rather what is remembered from trial to trial is affected. If such memory is lacking, one may not be able to form or enhance an explicit strategy that could contribute to the early phase of learning. Herzfeld et al. (2014) suggest that the error sensitivity depends on the history of past errors, which implies that the brain must store a memory of errors. Conversely, our results may suggest that this memory is affected in our patient group.

Other studies suggest that learning is constrained by the size of the error, rather than the sensitivity to error (Wei and Körding, 2009; Kim et al., 2018). The size of the error can be manipulated by the type of perturbation schedule that is used to elicit the adaptation. Using a gradual perturbation schedule in which participants never experience large errors, learning is suggested to be more implicit in nature (Orban de Xivry and Lefèvre, 2015) and may probe the slow process. Since the slow process learns only weakly from error, it may not depend, or depend less, on a memory of past errors, which would be consistent with the present results.

Our findings provide support for current disease management and clinical interventions that rely on the efficacy of errorless learning as an instructional method to facilitate learning in AD patients (Laffan et al., 2010; de Werd et al., 2013). That is, errorless learning is likely an intervention that capitalizes on the slow adaptive processes in these patients, and as we show here, these processes are still intact in early AD patients. It is unknown whether further progression of the disease affects the slow adaptive processes as well. The cerebellum is argued to play a central role in learning from small errors (Criscimagna-Hemminger et al., 2010; Vandevoorde and Orban de Xivry, 2019), but is typically not regarded as a key region in the etiology of early stage AD.

## Acknowledgments

This work was supported by a scholarship from the Donders Institute to K.S. W.P.M. is supported by the following grants: NWA-ORC-1292.19.298, NWO-SGW-406.21.GO.009 and Interreg NWE-RE:HOME.

## Supplementary material

**Supplementary Table 1:** MCMC diagnostics of the parameters. Parameter names in the table correspond to the parameters estimated by the model. For parameters that were estimated for individual participants (noted with an asterisk), the mean of the participants is listed. ESS - effective sample size; 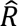 - convergence diagnostic; MCSE - Monte Carlo standard error.

| Parameter | AD patients |  |  | Control participants |  |  |
| --- | --- | --- | --- | --- | --- | --- |
| | ESS | $\hat{R}$ | MCSE | ESS | $\hat{R}$ | MCSE |
| $L_f - \mu$ | 1668.1 | 1.0025 | 0.00031 | 2538.0 | 1.0028 | 0.00008 |
| $L_f - \sigma$ | 2533.3 | 1.0017 | 0.00020 | 1276.8 | 1.0025 | 0.00012 |
| $L_s - \mu$ | 1341.9 | 1.0017 | 0.00005 | 468.4 | 1.0131 | 0.00008 |
| $L_s - \sigma$ | 2467.0 | 1.0012 | 0.00003 | 432.8 | 1.0125 | 0.00009 |
| $R_f - \mu$ | 562.6 | 1.0078 | 0.00228 | 650.4 | 1.0026 | 0.00037 |
| $R_f - \sigma$ | 420.5 | 1.0102 | 0.00227 | 336.3 | 1.0165 | 0.00056 |
| $R_s - \mu$ | 636.8 | 1.0054 | 0.00004 | 578.4 | 1.0039 | 0.00002 |
| $R_s - \sigma$ | 3013.7 | 1.0016 | 0.00001 | 874.7 | 1.0032 | 0.00001 |
| $\sigma_{state} - \mu$ | 432.9 | 1.0123 | 0.00022 | 536.1 | 1.0079 | 0.00015 |
| $\sigma_{state} - \sigma$ | 323.3 | 1.0153 | 0.00028 | 250.2 | 1.0214 | 0.00019 |
| $\sigma_{output} - \mu$ | 2226.7 | 1.0026 | 0.00032 | 1926.8 | 1.0016 | 0.00021 |
| $\sigma_{output} - \sigma$ | 2767.8 | 1.0022 | 0.00024 | 2238.6 | 1.0018 | 0.00016 |
| $L_f^*$ | 3074.4 | 1.0015 | 0.00041 | 2053.8 | 1.0018 | 0.00020 |
| $L_s^*$ | 4569.4 | 1.0014 | 0.00007 | 1184.7 | 1.0058 | 0.00010 |
| $R_f^*$ | 1068.6 | 1.0044 | 0.00284 | 849.2 | 1.0053 | 0.00064 |
| $R_s^*$ | 1403.7 | 1.0027 | 0.00004 | 1319.4 | 1.0020 | 0.00002 |
| $\sigma_{state}^*$ | 462.7 | 1.0137 | 0.00051 | 338.2 | 1.0209 | 0.00046 |
| $\sigma_{output}^*$ | 1008.5 | 1.0082 | 0.00072 | 791.9 | 1.0086 | 0.00055 |

**Supplementary Table 2:** Mean and 90% HDI of the model parameters. Parameter names in the table correspond to the parameters estimated by the model. For parameters that were estimated for individual participants (noted with an asterisk), the mean of the participants is listed.

|  | AD patients |  | Control participants |  |
| --- | --- | --- | --- | --- |
| Parameter | Mean | HDI | Mean | HDI |
| $L_f - \mu$ | 0.0700 | 0.0515 - 0.0922 | 0.0597 | 0.0530 - 0.0666 |
| $L_f - \sigma$ | 0.0391 | 0.0254 - 0.0577 | 0.0114 | 0.0045 - 0.0185 |
| $L_s - \mu$ | 0.0151 | 0.0120 - 0.0184 | 0.0187 | 0.0163 - 0.0216 |
| $L_s - \sigma$ | 0.0051 | 0.0026 - 0.0081 | 0.0032 | 0.0004 - 0.0064 |
| $R_f - \mu$ | 0.7765 | 0.6717 - 0.8453 | 0.9092 | 0.8927 - 0.9232 |
| $R_f - \sigma$ | 0.0878 | 0.0269 - 0.1760 | 0.0175 | 0.0024 - 0.0357 |
| $R_s - \mu$ | 0.9962 | 0.9946 - 0.9977 | 0.9972 | 0.9963 - 0.9980 |
| $R_s - \sigma$ | 0.0010 | 0.0001 - 0.0024 | 0.0006 | 0.0001 - 0.0014 |
| $\sigma_{state} - \mu$ | 0.0343 | 0.0273 - 0.0420 | 0.0239 | 0.0184 - 0.0298 |
| $\sigma_{state} - \sigma$ | 0.0150 | 0.0081 - 0.0240 | 0.0124 | 0.0080 - 0.0178 |
| $\sigma_{output} - \mu$ | 0.1076 | 0.0834 - 0.1323 | 0.0942 | 0.0788 - 0.1092 |
| $\sigma_{output} - \sigma$ | 0.0624 | 0.0445 - 0.0857 | 0.0385 | 0.0281 - 0.0518 |
| $L_f^*$ | 0.0700 | 0.0407 - 0.1082 | 0.0597 | 0.0455 - 0.0746 |
| $L_s^*$ | 0.0150 | 0.0088 - 0.0218 | 0.0187 | 0.0141 - 0.0242 |
| $R_f^*$ | 0.7752 | 0.6060 - 0.8859 | 0.9092 | 0.8782 - 0.9347 |
| $R_s^*$ | 0.9962 | 0.9937 - 0.9983 | 0.9972 | 0.9958 - 0.9984 |
| $\sigma_{state}^*$ | 0.0343 | 0.0207 - 0.0493 | 0.0239 | 0.0139 - 0.0353 |
| $\sigma_{output}^*$ | 0.1077 | 0.0813 - 0.1328 | 0.0942 | 0.0747 - 0.1126 |

## References

Albert MS, DeKosky ST, Dickson D, Dubois B, Feldman HH, Fox NC, Gamst A, Holtzman DM, Jagust WJ, Petersen RC, Snyder PJ, Carrillo MC, Thies B, Phelps CH (2011) The diagnosis of mild cognitive impairment due to Alzheimer’s disease: Recommendations from the National Institute on Aging-Alzheimer’s Association workgroups on diagnostic guidelines for Alzheimer’s disease. Alzheimers Dement 7:270–279.

Albert ST, Shadmehr R (2018) Estimating properties of the fast and slow adaptive processes during sensorimotor adaptation. J Neurophysiol 119:1367–1393.

Anguera JA, Reuter-Lorenz PA, Willingham DT, Seidler RD (2010) Contributions of Spatial Working Memory to Visuomotor Learning. J Cogn Neurosci 22:1917–1930.

Baddeley AD, Emslie H, Nimmo-Smith I (1994) Doors and People: A Test of Visual and Verbal Recall and Recognition. Thames Valley Test Company.

Braak H, Braak E (1991) Neuropathological stageing of Alzheimer-related changes. Acta Neuropathol 82:239–259.

Braak H, Braak E (1996) Development of Alzheimer-related neurofibrillary changes in the neocortex inversely recapitulates cortical myelogenesis. Acta Neuropathol 92:197–201.

Coltman SK, van Beers RJ, Medendorp WP, Gribble PL (2021) Sensitivity to error during visuomotor adaptation is similarly modulated by abrupt, gradual, and random perturbation schedules. J Neurophysiol 126:934–945.

Criscimagna-Hemminger SE, Bastian AJ, Shadmehr R (2010) Size of error affects cerebellar contributions to motor learning. J Neurophysiol 103:2275–2284.

de Werd MME, Boelen D, Rikkert MGMO, Kessels RPC (2013) Errorless learning of everyday tasks in people with dementia. Clin Interv Aging 8:1177–1190.

De Wit L, Kessels RPC, Kurasz AM, Amofa P, O’Shea D, Marsiske M, Chandler MJ, Piai V, Lambertus T, Smith GE (2022) Declarative Learning, Priming, and Procedural Learning Performances comparing Individuals with Amnestic Mild Cognitive Impairment, and Cognitively Unimpaired Older Adults. J Int Neuropsychol Soc 29:113–125.

De Wit L, Marsiske M, O’Shea D, Kessels RPC, Kurasz AM, DeFeis B, Schaefer N, Smith GE (2021) Procedural Learning in Individuals with Amnestic Mild Cognitive Impairment and Alzheimer’s Dementia: a Systematic Review and Meta-analysis. Neuropsychol Rev 31:103–114.

Dick MB, Andel R, Bricker J, Gorospe JB, Hsieh S, Dick-Muehlke C (2001) Dependence on Visual Feedback During Motor Skill Learning in Alzheimer’s Disease. Aging, Neuropsychology, and Cognition 8:120–136.

Fernandez-Ruiz J, Wong W, Armstrong IT, Flanagan JR (2011) Relation between reaction time and reach errors during visuomotor adaptation. Behav Brain Res 219:8–14.

Ferraro FR, Balota DA, Connor LT (1993) Implicit Memory and the Formation of New Associations in Nondemented Parkinson′s Disease Individuals and Individuals with Senile Dementia of the Alzheimer Type: A Serial Reaction Time (SRT) Investigation. Brain Cogn 21:163–180.

Ferrea E, Franke J, Morel P, Gail A (2022) Statistical determinants of visuomotor adaptation along different dimensions during naturalistic 3D reaches. Sci Rep 12:10198–10198.

Foundas AL, Leonard CM, Mahoney SM, Agee OF, Heilman KM (1997) Atrophy of the hippocampus, parietal cortex, and insula in Alzheimer’s disease: a volumetric magnetic resonance imaging study. Neuropsychiatry Neuropsychol Behav Neurol 10:81–89.

Haith AM, Huberdeau DM, Krakauer JW (2015) The influence of movement preparation time on the expression of visuomotor learning and savings. J Neurosci 35:5109–5117.

Herzfeld DJ, Vaswani PA, Marko MK, Shadmehr R (2014) A memory of errors in sensorimotor learning. Science 345:1349–1353.

Heuer H, Hegele M (2008) Adaptation to visuomotor rotations in younger and older adults. Psychol Aging 23:190–202.

Hong Y, Alvarado RL, Jog A, Greve DN, Salat DH (2020) Serial Reaction Time Task Performance in Older Adults with Neuropsychologically Defined Mild Cognitive Impairment. J Alzheimers Dis 74:491–500.

Howard IS, Ingram JN, Wolpert DM (2009) A modular planar robotic manipulandum with end-point torque control. J Neurosci Methods 181:199–211.

Hughes CP, Berg L, Danziger W, Coben LA, Martin RL (1982) A New Clinical Scale for the Staging of Dementia. Br J Psychiatry 140:566–572.

Hyman BT, Van Hoesen GW, Damasio AR, Barnes CL (1984) Alzheimer’s Disease: Cell-Specific Pathology Isolates the Hippocampal Formation. Science 225:1168–1170.

Inoue M, Uchimura M, Karibe A, O’Shea J, Rossetti Y, Kitazawa S (2015) Three timescales in prism adaptation. J Neurophysiol 113:328–338.

Julayanont P, Brousseau M, Chertkow H, Phillips N, Nasreddine ZS (2014) Montreal Cognitive Assessment Memory Index Score (MoCA-MIS) as a predictor of conversion from mild cognitive impairment to Alzheimer’s disease. J Am Geriatr Soc 62:679–684.

Keisler A, Shadmehr R (2010) A shared resource between declarative memory and motor memory. J Neurosci 30:14817–14823.

Kessels RPC, Remmerswaal M, Wilson BA (2011) Assessment of Nondeclarative Learning in Severe Alzheimer Dementia. Alzheimer Dis Assoc Disord 25:179–183.

Kim HE, Morehead JR, Parvin DE, Moazzezi R, Ivry RB (2018) Invariant errors reveal limitations in motor correction rather than constraints on error sensitivity. Commun Biol 1:19–19.

Kim S (2020) Bidirectional competitive interactions between motor memory and declarative memory during interleaved learning. Sci Rep 10:6916–6916.

Krakauer JW, Hadjiosif AM, Xu J, Wong AL, Haith AM (2019) Motor Learning. Compr Physiol 9:613–663.

Kruschke JK (2015) Doing Bayesian Data Analysis, 2nd ed. Elsevier.

Kruschke JK (2018) Rejecting or Accepting Parameter Values in Bayesian Estimation. Advances in Methods and Practices in Psychological Science 1:270–280.

Laffan AJ, Metzler-Baddeley C, Walker I, Jones RW (2010) Making errorless learning more active: self-generation in an error free learning context is superior to standard errorless learning of face-name associations in people with Alzheimer’s disease. Neuropsychol Rehabil 20:197–211.

Lee J-Y, Schweighofer N (2009) Dual adaptation supports a parallel architecture of motor memory. J Neurosci 29:10396–10404.

Leow L-A, Gunn R, Marinovic W, Carroll TJ (2017) Estimating the implicit component of visuomotor rotation learning by constraining movement preparation time. J Neurophysiol 118:666–676.

Malfait N, Ostry DJ (2004) Is interlimb transfer of force-field adaptation a cognitive response to the sudden introduction of load? J Neurosci 24:8084–8089.

Mazzoni P, Krakauer JW (2006) An implicit plan overrides an explicit strategy during visuomotor adaptation. J Neurosci 26:3642–3645.

McDougle SD, Wilterson SA, Turk-Browne NB, Taylor JA (2022) Revisiting the Role of the Medial Temporal Lobe in Motor Learning. J Cogn Neurosci 34:532–549.

McKhann GM, Knopman DS, Chertkow H, Hyman BT, Jack CR Jr, Kawas CH, Klunk WE, Koroshetz WJ, Manly JJ, Mayeux R, Mohs RC, Morris JC, Rossor MN, Scheltens P, Carrillo MC, Thies B, Weintraub S, Phelps CH (2011) The diagnosis of dementia due to Alzheimer’s disease: recommendations from the National Institute on Aging-Alzheimer’s Association workgroups on diagnostic guidelines for Alzheimer’s disease. Alzheimers Dement 7:263–269.

Nasreddine ZS, Phillips NA, Bédirian V, Charbonneau S, Whitehead V, Collin I, Cummings JL, Chertkow H (2005) The Montreal Cognitive Assessment, MoCA: a brief screening tool for mild cognitive impairment. J Am Geriatr Soc 53:695–699.

Oldfield RC (1971) The assessment and analysis of handedness: the Edinburgh inventory. Neuropsychologia 9:97–113.

Orban de Xivry J-J, Lefèvre P (2015) Formation of model-free motor memories during motor adaptation depends on perturbation schedule. J Neurophysiol 113:2733–2741.

Rouleau I, Salmon DP, Vrbancic M (2002) Learning, retention and generalization of a mirror tracing skill in Alzheimer’s disease. J Clin Exp Neuropsychol 24:239–250.

Sarwary AME, Wischnewski M, Schutter DJLG, Selen LPJ, Medendorp WP (2018) Corticospinal correlates of fast and slow adaptive processes in motor learning. J Neurophysiol 120:2011–2019.

Schmand BA, Lindeboom J, Harskamp FV (1992) NLV: Nederlandse leestest voor volwassenen: handleiding. Swets & Zeitlinger.

Shadmehr R, Brandt J, Corkin S (1998) Time-Dependent Motor Memory Processes in Amnesic Subjects. J Neurophysiol 80:1590–1597.

Smith GE, Petersen RC, Parisi JE, Ivnik RJ, Kokmen E, Tangalos EG, Waring S (1996) Definition, course, and outcome of mild cognitive impairment. Aging, Neuropsychology, and Cognition 3:141–147.

Smith MA, Ghazizadeh A, Shadmehr R (2006) Interacting adaptive processes with different timescales underlie short-term motor learning. PLoS Biol 4:e179.

Taylor JA, Ivry RB (2011) Flexible cognitive strategies during motor learning. PLoS Comput Biol 7:e1001096.

Taylor JA, Krakauer JW, Ivry RB (2014) Explicit and implicit contributions to learning in a sensorimotor adaptation task. J Neurosci 34:3023–3032.

Trewartha KM, Garcia A, Wolpert DM, Flanagan XJR, Flanagan JR (2014) Fast but fleeting: adaptive motor learning processes associated with aging and cognitive decline. J Neurosci 34:13411–13421.

van Halteren-van Tilborg IADA, Scherder EJA, Hulstijn W (2007) Motor-skill learning in Alzheimer’s disease: a review with an eye to the clinical practice. Neuropsychol Rev 17:203–212.

Van Hoesen GW, Hyman BT, Damasio AR (1991) Entorhinal cortex pathology in Alzheimer’s disease. Hippocampus 1:1–8.

van Tilborg IADA, Hulstijn W (2010) Implicit Motor Learning in Patients with Parkinson’s and Alzheimer’s Disease: Differences in Learning Abilities? Motor Control 14:344–361.

Vandevoorde K, Orban de Xivry J-J (2019) Internal model recalibration does not deteriorate with age while motor adaptation does. Neurobiology of Aging 80:138–153.

Verhage F (1964) Intelligence and Age: Study with Dutch People Aged 12-77 (in Dutch). Assen: Van Gorcum.

Wei K, Körding K (2009) Relevance of error: what drives motor adaptation? J Neurophysiol 101:655–664.

Willingham DB, Peterson EW, Manning C, Brashear HR (1997) Patients with Alzheimer’s disease who cannot perform some motor skills show normal learning of other motor skills. Neuropsychology 11:261–271.

Wolpe N, Ingram JN, Tsvetanov KA, Henson RN, Wolpert DM, Cam-CAN, Rowe JB (2020) Age-related reduction in motor adaptation: brain structural correlates and the role of explicit memory. Neurobiol Aging 90:13–23.

Wong S, Irish M, Savage G, Hodges JR, Piguet O, Hornberger M (2019) Strategic value-directed learning and memory in Alzheimer’s disease and behavioural-variant frontotemporal dementia. J Neuropsychol 13:328–353.

Zanetti O, Zanieri G, Giovanni GD, De Vreese LP, Pezzini A, Metitieri T, Trabucchi M (2001) Effectiveness of procedural memory stimulation in mild Alzheimer’s disease patients: A controlled study. Neuropsychological Rehabilitation 11:263–272.

